# Design and development of a vortex ring generator to study the impact of the ring as a gust

**DOI:** 10.1101/2020.10.12.331777

**Authors:** Dipendra Gupta, Sanjay P. Sane, Jaywant H. Arakeri

## Abstract

We present a simple method to generate discrete aerodynamic gust under controlled laboratory condition in the form of a vortex ring which, unlike conventional methods of perturbation, is well studied and highly controllable. We characterized the flow properties of the vortex ring using flow visualization and novel light bead method. Reynolds number of the vortex ring, based on its average propagation velocity and nozzle exit diameter, was 16000. We demonstrate this method by studying the impact of head-on gust on freely flying soldier flies, Reynolds number of which, based on its wingtip velocity and mean wing chord, was 1100. We also present simple theoretical models to characterize the vortex ring based on generating conditions. The device can also be used to generate continuous gust in any direction and can be applied, in general, to study the gust response of natural fliers and swimmers, man-made micro aerial vehicles and aquatic plant lives.

## INTRODUCTION

The environment in which biological organisms locomote is characterized by the preponderance of turbulence, that comprises of discrete gusts and random continuous turbulence (Etkin 1981). Discrete gusts, that refer to a sudden and sharp change in wind speed, are encountered in the wakes of large objects or at the edge of convective disturbances while the latter can be referred to as a chaotic motion of atmospheric wind and is usually described using a statistical approach. Nonetheless, they are able to forage successfully in these extreme environmental conditions, which necessitates the study of their response and recovery strategies in such conditions for better design of highly efficient unmanned autonomous vehicles (UAVs). But, even before we do that, one needs to generate a well-characterized gust. Some methods of generation of turbulence and gust reported in the existing literature, in context of natural fliers, include grid generated turbulence (Combes & Dudley 2009; Crall et al. 2017; Ravi et al. 2015), von-Karman vortices (Ravi et al. 2013; Ortega-Jimenez et al. 2013; Ravi et al. 2016; Engels et al. 2016; Jakobi et al. 2018; Ortega-Jimenez et al. 2014), compressed air jet (Vance et al. 2013; Boerma et al. 2019) and Helmholtz coils (Ristroph et al. 2010; Ristroph et al. 2013; Beatus et al. 2015). However, one of the main problems associated with such studies is that of spatio-temporal characterization of the gust. Even, some of these methods of generating the perturbation in these experiments are not realistic in nature, thus, entailing a new method of gust generation that can overcome these limitations.

We here present a simple device to generate a discrete head-on gust in form of a vortex ring. An impulsively started flow around sharp edges when separates can lead to vortex formation that rolls up into a spiral form, finally giving rise to a vortex ring, that propagates with its own self-induced velocity due to vorticity concentrated in the core region. One interesting feature of the vortex ring, that makes it a suitable candidate for its employability as a gust, is its high impulse and presence of vorticity mostly in its core. A recent study also employed this method to perturb live freely swimming fish in water (Seth et al. 2017). It was, however, focused just on the impact of the ring to observe a distinct change in fin motion, rather than characterizing its flow properties and the response of the fish due to perturbation, and hence, necessitating a detailed study of the evolution of the ring with space and time that can help better understand the response of these natural organisms under such severe environment. To facilitate the reader in designing a well-characterized gust generator, we first provide a recommendation, based on simple theoretical models, for estimation of the flow properties of the ring based on generating parameters. We then discuss the experimental setup for this method and the spatio-temporal characterization of the flow properties of the gust generated by this method, and finally discuss, in brief, the application of this method as a gust to study its impact on insect flight.

## METHODS AND MATERIALS

### Theoretical models

Parameters like diameter, propagation speed and circulation of the vortex ring are some of the important properties to consider for its characterization. Considering its application as a gust, we discuss here only its diameter and propagation speed. The diameter of the rings depends on the exit diameter of the cylindrical tube (D_0_) and the distance piston is moved (Auerbach 1988) (called here as stroke length) and can be equal to, smaller or larger than the exit diameter of the tube (Sullivan et al. 2008). During the formation process (i.e. time it takes to form a fully developed ring), the ring entrains the surrounding fluid as it propagates, and gives rise to a bubble-like structure called vortex bubble (D_vb_), the diameter of which is clearly larger than that of the ring (D_r_). Even after the formation process, the entrainment can occur followed by subsequent detrainment, the balance between which precludes substantial change in its diameter (Maxworthy 1972).

The momentum (or more precisely impulse) of the ring is determined by the momentum imparted to the surrounding fluid in the tube by the piston. It depends on the type of motion imposed on the piston, piston travel time, viscosity, and its radius and circulation. During the formation process, the ring accelerates, and then may gain constant velocity or slow down. The slowing down may be attributed to the entrainment of surrounding fluid (Reynolds 1876), entrainment followed by detrainment (Maxworthy 1972), viscous diffusion of the core (Saffman 1970) or vortex instabilities (Glezer & Coles 1990). It is worth noting that the turbulent vortex ring, in compared to the laminar one, is characterized by a rapid growth rate of its diameter and shedding of significant vorticity to the wake, and consequently, rapid decrement in its propagation velocity.

It is now evident that prior knowledge of piston movement time (called here as stroke time, T_p_) and slug length (L) can help the experimentalists predict the flow properties of the ring. We, here, provide general working relations based on simple conservation laws to achieve this objective.

#### A. Estimation of slug length (L)

Some volume of fluid gets ejected out of the nozzle due to impulsive motion of the piston, the effective length of which is called slug length. Based on the velocity profile and the time of piston motion, one can estimate the slug length as:

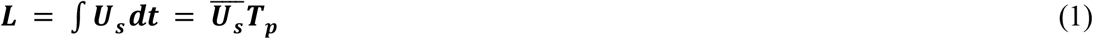

Where 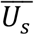 is the average uniform velocity of the slug after it is fully formed into a vortex ring. (1) can be also be derived using volume conservation and continuity equations. Because the volume of fluid coming out of the nozzle is equal to that pushed by the piston (Figure 1A) (Das et al. 2017), the slug length can be given as

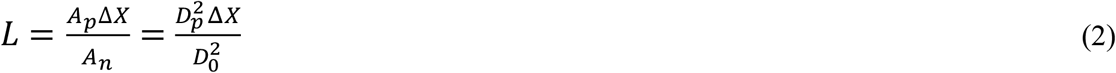

**Figure 1:**
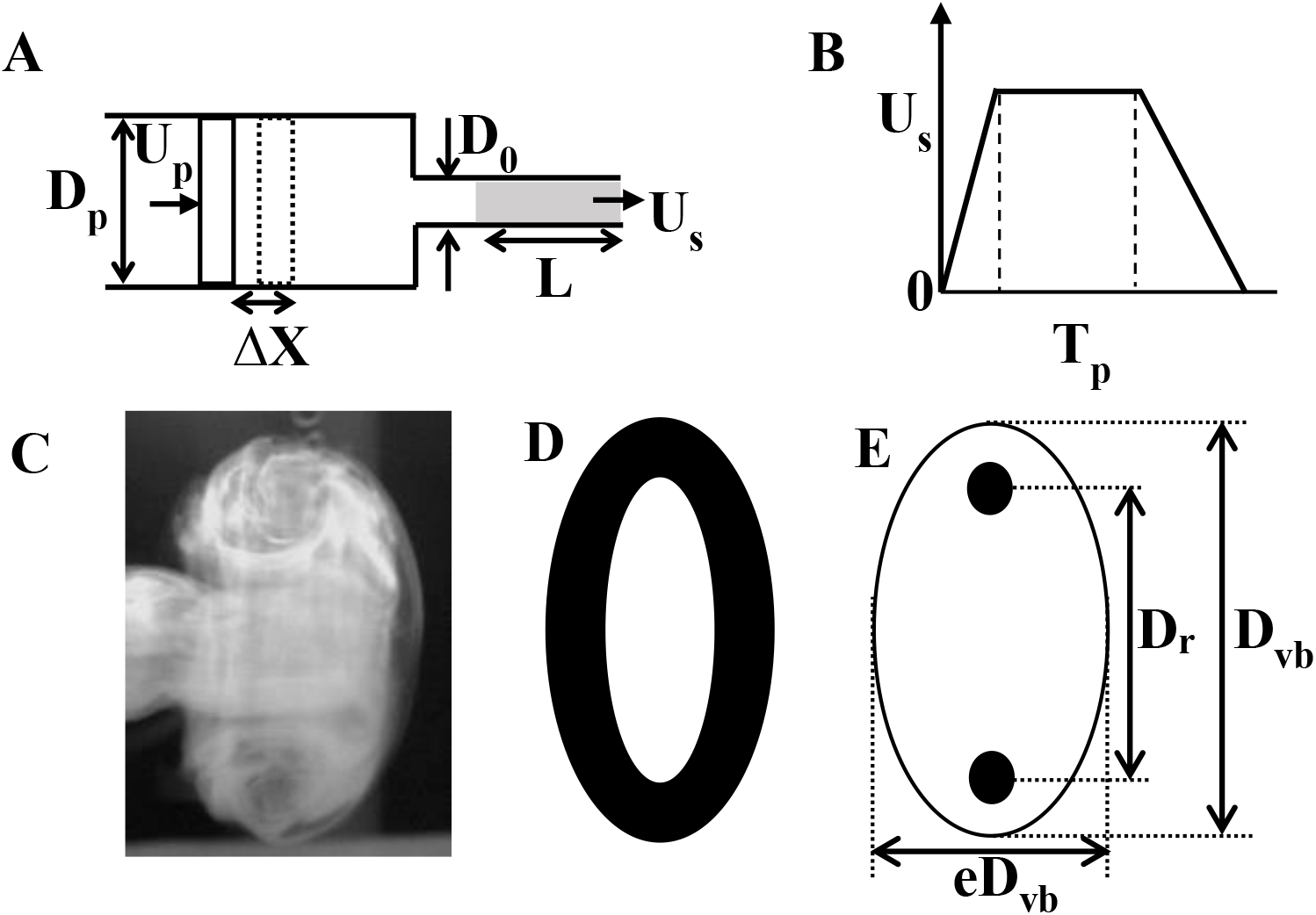
Geometric details of Vortex ring, and ring generator. (A) Piston-nozzle arrangement for generating vortex ring. Dashed line is the outline of piston after it has moved by ΔX distance and grey fill is the length of fluid that comes out of the nozzle and is called here as slug length (L). (B) Variation of piston velocity/slug velocity with time. D_p_ and U_p_ are diameter and velocity of piston respectively, *D*_0_ and U_s_ are exit diameter and exit velocity of the nozzle respectively. T_p_ is the total time for which piston moves. (C) Side view, (D) isometric view and (E) assumed line diagram of side view of the ring. Highly dark circle in E denotes the core of the vortex ring, where most of the vorticities are concentrated. D_r_, D_vb_ and e denote diameter of the ring (distance between the cores), diameter of the vortex bubble including entrained air and eccentricity of the ellipsoid respectively.

Where A_p_ and A_n_ are area of piston and nozzle respectively.

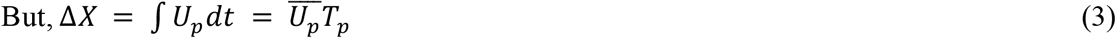

Where 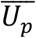 is average velocity of the piston.

Applying continuity equation on piston and nozzle exit,

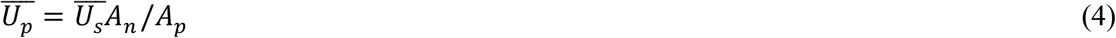

From (1), (2) and (3), 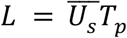

Alternatively, one can estimate L using vortex ring dimension. For this, we may assume that vortex ring takes the form of an ellipsoid with eccentricity ‘e’ which is the ratio of minor to major axes, and since volume of fluid coming out of nozzle (Ω_n_) is equal to that contained in the ring (Ω_r_) (Figure 1C-E) (Sullivan et al. 2008).

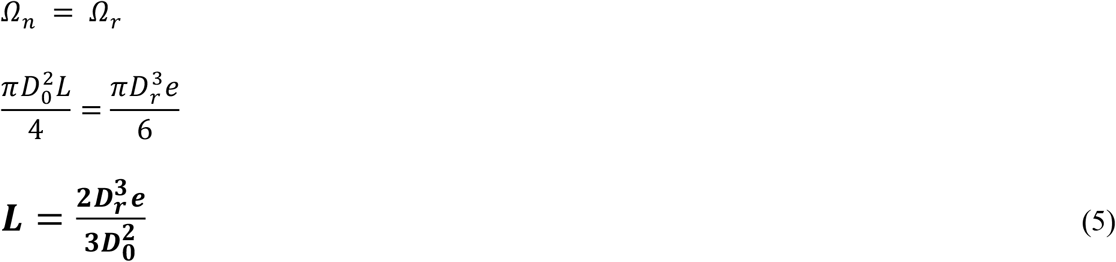

We here note that the volume of the ring considered here doesn’t take into account entrainment, and hence, doesn’t represent the actual volume. Nonetheless, (5) holds good experimentally (Sullivan et al. 2008). In fact, due to entrainment, the total mass, and hence, the total volume of the ring increases. The volume of the vortex bubble is given as (Ω_vb_)= Ω_n_ + Ω_entrained_, and Ω_entrained_ is about (20-40)% total volume of the ring (Auerbach 1991; Dabiri & Gharib 2004).

#### B. Estimation of core radius (a)

For small time and thin vortex core, the core radius of the ring can be estimated as (Saffman (1970):

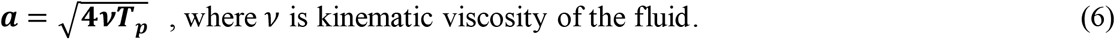

#### C Estimation of slug velociy 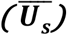

The velocity of the fluid ejected out of the nozzle after it is fully formed into a ring can be estimated as (Sullivan et al. 2008)

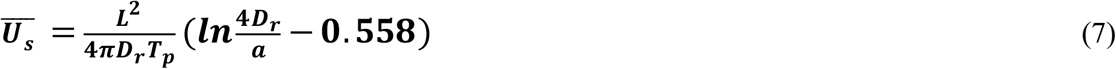

where L^2^/2T_p_ gives the estimation of circulation of the slug.

#### D. Estimation of translational velocity (U_r_)

Following momentum conservation, the momentum of the slug ejected out of nozzle can be equated to the momentum of the vortex bubble, i.e.

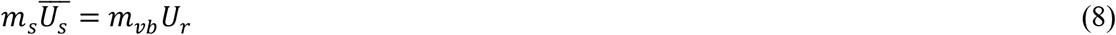

where m_s_ is the mass of fluid displaced by the piston, and mvb and U_ravg_ are the mass and the velocity of the ring with entrained air (i.e. vortex bubble).

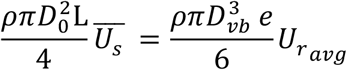

Using (1) and solving,

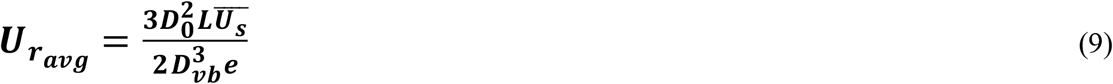

Note that D_vb_ is the diameter of the ring including both slug and entrained fluid. The piston velocity can be determined from (1) or (7).

### Experimental setup

The experimental set-up consists of a 60 cm long, 30cm square cross-section dismountable clear Perspex chamber, a 40 cm long, 3.76 cm internal diameter (D_0_) cylindrical PVC nozzle (2mm thick), a 12-inch 100W and 8 Ω speaker), a digital to analog converter NI-DAQ, a high voltage high current direct coupled (DC) amplifier and a high-speed camera (Miro EX4) (Figure 2A).

**Figure 2:**
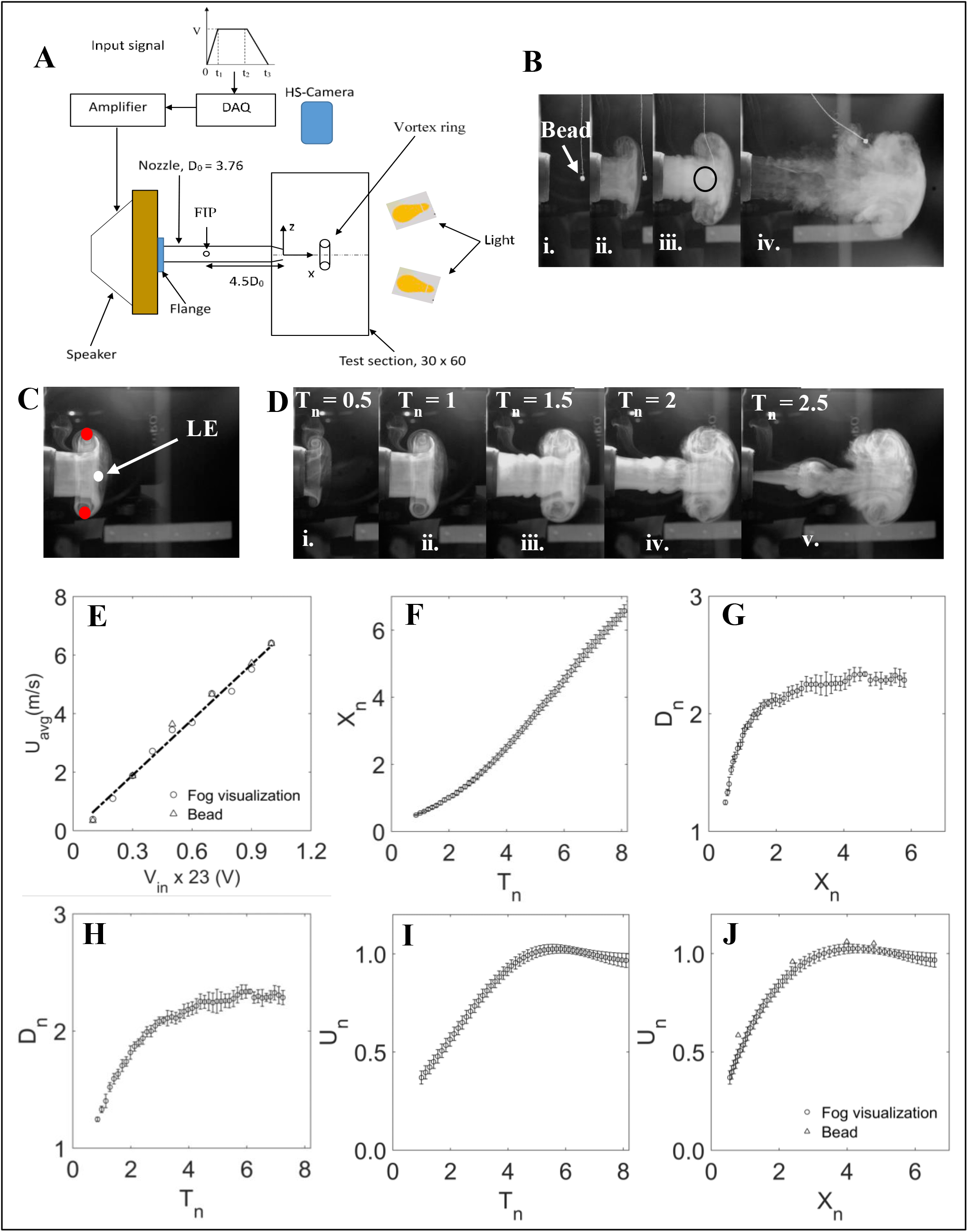
Generation and characterization of vortex ring. (A) Different components of the experimental setup and their arrangement. Test chamber, Speaker, nozzle, Infrared (IR) motion sensor, Fly releasing dust (FRD), halogen lamp, cameras, DAC and amplifier together constitute the gust generator system. ROI is region of interest where the encounter of insect with the gust is desired. Left top shows input signal to DAC with voltage amplitude V, rising time, t1, constant duration t_2_-t_1_ and fall time t_3_-t_2_-t_1_. This signal is converted into analog form, amplified by an amplifier and fed to the speaker for vortex ring generation. Fog particles were injected into nozzle through fog injection port (FIP) and high-speed camera (HS-Camera) was placed horizontal to record the lateral view of the vortex ring as it propagates. All dimensions are in cm. (B-J) Flow visualization and characterization of vortex ring. (B) Effect of gust (U_avg_ = 6.4 m/s) on a freely hanging Styrofoam bead. Position of bead (i) when there is no gust, (ii) just before the gust, (iii) during gust, and (iv) after gust. The bead translates with the gust as seen in (iii) unless it reaches the position where it can no longer be carried by gust due to finite length of the thread. Black circle in (iii) shows the position of the bead when it is at the centre of the vortex ring. (C) White and red circle denote leading edge (LE) and extreme end of the ring, that were tracked to calculate its axial position and diameter respectively. (D) Flow visualization at different time instances for U_avg_ = 1.9 m/s. The ring propagates from left to right along X in each figure. T_n_=0 indicates the time instance when no ring is formed. (E) Average propagation velocity of the ring as a function of voltage input to the speaker. Velocity of the ring obtained using fog visualization (circles) and bead method (triangles) are in good agreement for different input voltages. Dashed line is *U_avg_* = 0.2745 *V_in_*, R^2^ = 0.99. (F)-(J) Non-dimensional flow properties of vortex ring with U_avg_=6.4 m/s and vortex bubble diameter 8.6 cm measured using flow visualization and bead method. X_n_=0 indicates the centre of nozzle exit. (F) Position of ring with time. (G-H) Diameter with axial distance (G), and with time (H). (I-J) Propagation velocity showing time independency after T_n_=4 (I) and uniformity after X_n_= 3 from nozzle exit (J). Values are mean ± SD. See text for non-dimensional parameters.

The large Perspex chamber served as a closed test section where we generated vortex ring. The large dimension of the test chamber compared to the diameter of the vortex ring reduces any effects of external air currents on experimental observations, and hence, aids in maintaining a still ambient fluid (Das et al. 2017). We carried the experiments in a closed room with a controlled level of humidity and an ambient temperature of 20°C. The speaker was enclosed in a 40cm x 40cm x 5cm wooden chamber (driving section) on the diaphragm side, and each side of the chamber was glued with fevicol and pin hammered to ensure airtightness. A 5cm diameter hole was cut at the centre of the 40cm x 40cm face of the wooden chamber to facilitate attachment of the nozzle via a PVC flange. A rubber gasket was used in between flange and wooden chamber to eliminate any air leakage. The nozzle was sharp chamfered by an angle of 9° at the exit and smooth chamfered at entry. The speaker attached with the nozzle was then fitted to the test chamber through a 4cm hole cut on its longest side as shown in Figure 2A.

### Input Signal

We initially synthesized a trapezoidal signal (Figure 2A) using NI-LabVIEW and stored it in computer memory. The signal consisted of three parts: acceleration (i.e. rise), constant velocity, and deceleration (i.e. falling). We chose the time period for each portion of the signal (t_1_= 100 μs, t_2_- t_1_ = 30 ms and t_3_- t_2_- t_1_=100 ms) such that it resulted desired velocity of the vortex ring. Such a large deceleration time eliminates the formation of opposite sign stopping vortex (Das et al. 2017) which is formed, on abrupt stopping of piston, due to separation and rolling of the secondary boundary layer formed on the outer surface of the tube due to induced velocity of primary vortex ring (Didden 1979; Lim & Nickels 1995; Pullin & Perry 1980; Weigand & Gharib 1997; Shariff & Leonard 1992).

We converted the signal into analog form for physical output using NI −c9263, amplified it using an in-house DC power amplifier, and finally fed it to the speaker, resulting in the formation of a vortex ring at the exit of the nozzle.

### Formation of vortex ring

The signal, when fed to the speaker, displaces its diaphragm impulsively by some distance in the wooden chamber. The diaphragm, in turn, imparts its momentum to the surrounding air, thus causing an equivalent volume of air being pushed out from the chamber into the nozzle, generating a layer of vorticity (boundary layer) in the inner wall of the nozzle to satisfy a no-slip condition. As the high-speed slug of air comes out of the nozzle, this boundary layer forms a cylindrical vortex sheet that immediately starts rolling up into a spiral form, giving rise to a vortex ring (Lim & Nickels 1995).

The present experimental set-up, unlike conventional piston-cylinder configuration for vortex ring generation (Didden 1979; Lim & Nickels 1995; Pullin & Perry 1980; Weigand & Gharib 1997; Kumar et al. 1995; Maxworthy 1974; Auerbach 1991; Irdmusa & Garris 1987; Glezer 1988; Gharib et al. 1998; Glezer & Coles 1990; Cater et al. 2004; Sullivan et al. 2008; Maxworthy 1977; Auerbach 1987) has speaker diaphragm (analogous to piston) in the driving section far ahead of the nozzle, thus eliminating the generation of piston vortex (Das et al. 2017). Piston vortex (Allen & Auvity 2002; Glezer 1988; Glezer & Coles 1990; Cater et al. 2004) is formed in front of the piston as it moves ahead in the tube, due to removal of a boundary layer of the tube wall as the piston advances forward (Allen & Auvity 2002). Further, the long nozzle (11 times its diameter) forces the slug of fluid coming out only from the front part of the nozzle, not from the entry region where the nozzle is attached to the wooden chamber, thereby ensuring the elimination of the possibility of having any disturbances similar to the orifice generated vortex ring (Das et al. 2017).

### Characterization of gust

We characterized the gust using two different techniques: fog visualization and Styrofoam bead method. Both the methods were carried out independently, and flow properties were characterized for each method.

### Fog visualization

We used Antari fog particles as seeding particles for visualization of the vortex ring. The average particle size of the fog was of the order of 1-2 μm. The fog was first filled into a 500 ml wash bottle and injected through a fog injection port (FIP) on the nozzle (Figure 2A). The port was made on the upper circumference of the nozzle, at 18 cm (4.5D_0_) away from its exit plane. The circumferential (lower, upper or sidewise) position of the port didn’t seem to affect the visualization. Its longitudinal position, however, determined whether any fog particles would be present at the exit of the nozzle before ring formation (i.e. background fog at nozzle exit). The choice of its longitudinal position (4.5D_0_) eliminated the possibility of any background fog formation at the nozzle exit.

To prevent fog accumulation inside the test chamber, a 5cm square window was hinged on the longer side opposite to nozzle. We kept the window closed during visualization to eliminate any effects of external air current on the ring propagation and its trajectory. It was opened only after the completion of the recording. This ensured that there remained no fog inside the chamber before the start of the next trial.

Figure 2D shows the formation of the vortex ring at 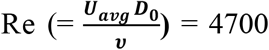 at different time instances, where *U_avg_* is average propagation speed of the vortex ring (defined later), *D*_0_ is nozzle exit diameter and *ν* is kinematic viscosity. It can be observed that vortex sheet emanates from the edge of the nozzle and rolls up to form a spiral, as the slug of fluid gets ejected through the nozzle. The vortex ring continues to propagate while drawing more fluids from the surrounding ambient. As discussed above, we also don’t observe any evidence of secondary and piston vortices. In this sense, the vortex ring generated here is clean and free from the effect of these disturbances.

### Styrofoam Bead method

We experimentally found the density of Styrofoam bead (6.52 kg/ m^3^) to be of an order of the density of air (see table S1), and so, we expected that the velocity of the bead, on an encounter with gust, should be of the nearly same magnitude as that of the gust. With this expectation, we employed this novel method to determine the velocity of gust at different axial locations. For which we suspended a Styrofoam bead using thin sewing thread from the test chamber ceiling, such that it rests on the centreline of the nozzle exit, and hit it by the vortex ring. The gust imparts an impulsive motion to the bead as it propagates past the bead. We repeated this process by hanging bead at different axial locations on the centreline of nozzle exit to find the velocity of the bead, and hence, the gust at that axial locations. One such trial for U_avg_ = 6.4 m/s is shown in Figure 2B.

We used a 12-bit CMOS camera (Phantom Miro EX4) fitted with Nikon 18-70 mm focal length lens to record the flow images for both these methods at 1200 fps and 50 μs exposure time. To take care of such a low exposure time, we illuminated the background using two 1000W halogen lamps. We placed the camera horizontal such that a lateral view of vortex ring propagation would be captured in a plane perpendicular and vertical to the nozzle exit plane. The external diameter served as a calibration scale for the images.

We defined non-dimensional time based on exit diameter of nozzle and ring average velocity, time **(t)** as 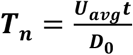. Similarly, axial distance from nozzle exit was non-dimensionalized with the exit diameter and is given by 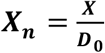. The dimensionless diameter of the nozzle is given by ***D_n_ = D_vb_/D_0_***, where ***D_vb_*** is the instantaneous diameter of vortex bubble (i.e. diameter of the ring with entrained air), and dimensionless velocity of the ring is given by 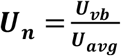 where ***U_vb_*** is instantaneous velocity of the vortex bubble.

## RESULTS AND DISCUSSION

Considering the aim of its application as a discrete gust, we here, focus on its spatial and temporal evolution of vortex ring and its translational velocity. We observed that the ring velocity varies linearly from 0.4 m/s to 6.4 m/s for input voltages ranging from 2.3 V to 23 V (*U_avg_* = 0.2745*V_in_, R*^2^ = 0.99) (Figure 2E), the corresponding Re for which ranges from 1 x 10^3^ to 1.6 x 10^4^. Such strong dependence of ring’s translational velocity on voltage shows that simply modulating the amplitude of input signal, one can achieve vortex ring of different strength, thus allowing high controllability over its flow properties. The bead velocity and the average velocity of gust measured using flow visualization matches very well for each voltage input, thus validating our hypothesis that the bead can be used as a tracer particle in air for such flow. We note that the velocity changes in a similar fashion with time and axial space for each of the voltage input, and as a special case, we also discuss here the ring flow properties for Re = 16000 (U_avg_=6.4 m/s) (Figure 2F-J).

The ring propagates quadratically with time up to T_n_ =4 after which it moves linearly until it reaches near the wall of the test chamber where wall effects can be expected to be dominant (Figure 2F). Because the core of the ring was not visible in all images and across all videos, we tracked the point LE to measure its axial position, and lateral extremes of vortex ring to measure its diameter (Figure 2C). The diameter of the ring also grows quadratically with space from 1.25D_0_ to 2.25D_0_ until X_n_=3 (Figure 2G), after which it attains space invariant final size of 2.3 ± 0.03 D_0_. We note that the diameter measured here is the diameter of the vortex bubble and not the ring. Because entrained fluid generally constitutes about 20-40% of the total volume of fluid carried by tube generated vortex ring (Auerbach 1991; Dabiri & Gharib 2004) and for the present study, the eccentricity, e=0.62, if we assume entrained mass fraction is 30%, it yields D_r_ = 0.89D_vb_.

Further, the diameter first increases and then becomes constant with time, and can be fitted with a quadratic equation with time for the entire observation period (Figure 2H). The correlation equations for these flow properties with time and axial distance are given in equations (1–10). For its application as a gust, its diameter should be chosen such that the characteristic length of the subject is well within the circumference. For instance, the diameter, in case of birds, bats and insects, should be at least equal to their wingspan, and for fish, it should be at least equal to the body length which response is to be studied, so that, under controlled laboratory setup, their flight and swimming can be contained within the ring. It is, however intuitively, suggested that the ring diameter at least twice the characteristic length should be preferred to ease a greater degree of freedom in their flight. While it can be increased by more than twice, one should also bear in mind that it can limit the maximum gust velocity (the smaller the ring, the faster it travels (Saffman 1970)) in addition to posing experimental difficulties in case of large animals.

The non-dimensional ring velocity, calculated by applying second-order central difference scheme to axial position, becomes time-invariant after T_n_ = 4 (Figure 2I) and uniform after Xn = 3 from the nozzle exit (Figure 2J). The absolute value of the average speed of the ring for Re=16000 is 6.4 m/s. The representative instantaneous streamlines of the ring and its axial and radial velocity distribution at a different location and different time instances are given in Figure S1 and S2. We note that the average velocity of the ring was obtained by averaging the velocity after X_n_ = 3 after which it becomes nearly constant, and for the bead method, the maximum velocity of bead, which it attains in the first 5 frames of the image sequences, was considered for comparison with the average velocity obtained using fog visualization (Figure 2E,J). Any motion of the bead is due to the impulsive force of the ring, and hence, consideration of the maximum velocity is well justified. For the application of the ring as a gust, its translational velocity should be of the order of forward velocity of the subject under consideration. While it may hold for birds, bats and fish, in case of insects that flap their wings at higher rates, it may not induce any changes in its body kinematics (as was evidenced in our experiments as well), and may require considerably higher velocity, of the order of wingtip velocity. Based on our experimental result, we found the following relations applicable to the vortex ring at U_avg_=6.4 m/s.

### 1. Axial position

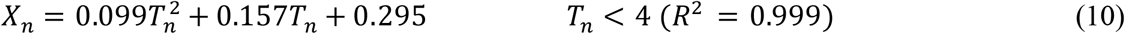

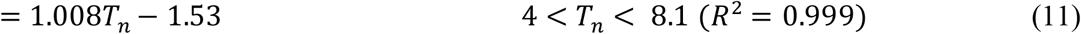

### 2. Diameter

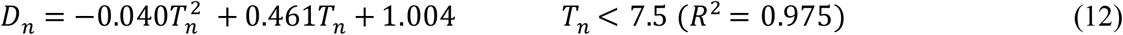

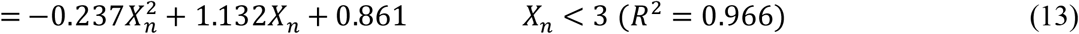

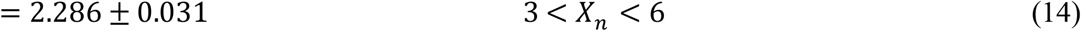

### 3. Translational velocity

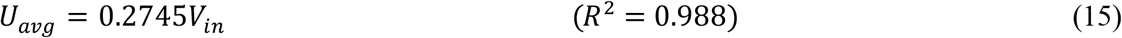

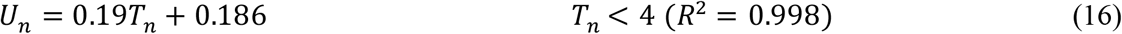

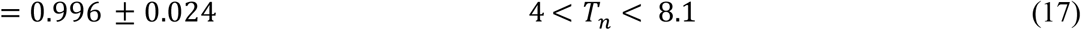

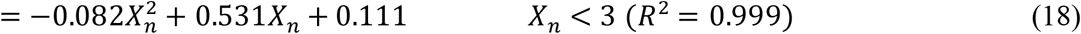

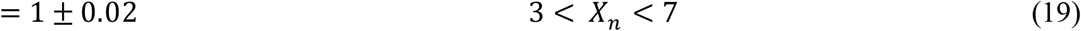

We note that the maximum standard deviation in ring properties measured for three trials was less than 10% and average standard deviation less than 5% of their mean values, implying high repeatability of the measured values.

#### Validation of the theoretical model

The value of the diameter of the ring without considering entrainment (Dr) as obtained above gives L=12.83 cm (eqn 5) which when substituted (together with Tp =30.1 ms) in eqn 1 results the slug velocity, Us of 4.26 m/s. Similarly, the slug velocity calculated using core radius based on T_p_ (a=1.35 mm here, see eqn (6)) is 2.8 m/s (eqn 7). We note that the time of piston motion during deceleration is neglected as this time is assumed not to contribute to the formation of the ring. These predicted values of slug velocity, after considering entrainment (eqn 9), give the average velocity of the ring as ~3 m/s and ~2m/s respectively. The estimated ring velocity is, however, lower than the experimental value (6.4 m/s). The exact reason for this discrepancy is not known, but we believe that this error may be mainly due to error in consideration of the particular time of the piston motion as, due to experimental limitation, we could not measure the exact time for which the piston (speaker diaphragm) moves. Further, the diaphragm does not get displaced uniformly around the circumference of the speaker since it is fixed to the speaker circumferentially and displaces mostly in the central region. Despite these limitations, the estimated value is of the order of the experimental values and hence, eqns 1-9 can be used to estimate the ring velocity. It is also interesting to note that the estimated L gives L/D_0_ = 3.41 which is very close to the universal formation time (L/D0 ~ 4) for vortex ring at which it attains maximum circulation and impulse, and after which, the transition from an isolated form to that followed by trailing jet occurs (Gharib et al. 1998). However, we can still see some trailing jet in our experiments (Figure 2B). We assumed that the velocity of the trailing jet is minimal compared to that of the ring and, due to viscous diffusion, it does not last longer (Lim & Nickels 1995).

#### Application of vortex ring as a gust

To show the effectiveness of the vortex ring as a discrete head-on gust, we present here some results on change in flight kinematics of four flies when subjected to the gust.

We selected vortex ring velocity of 6.4 m/s to perturb the free-flying soldier flies and recorded their flight motion at 4000 FPS using two synchronized, high-speed cameras (Phantom VEO 640L and Phantom V611) for more than 80 trials, 14 of which we calibrated and digitized using MATLAB based routines-easywand5 (Theriault et al. 2014) and DLTdv7 (Hedrick 2008) respectively to study some general trends in change in their body and wing kinematics due to gust. We first down-sampled the image data to 1000 frames per second, and then digitized and tracked head, abdomen, each wing base and tip to get their 3D position in the global reference frame. We assumed Centre of Mass (CoM) at one-third body length from the abdomen, and used it to represent the flight trajectory. X, Y, and Z represent the axial, lateral, and vertical axes of the flies respectively.

We calculated total velocity (forward speed or velocity magnitude) as the resultant of three velocity components. For this, we first applied fourth-order low pass Butter-worth filter with cut-off frequency 200Hz to the corresponding CoM data to minimize any digitization error, and then calculated the velocity along each axis using second-order central difference scheme. We non-dimensionalized velocity by the product of the body length of the flies and their wingbeat frequency (f). Similarly, we calculated the body roll angle (γ) with respect to the horizontal plane as the elevation angle of the vector joining the wing base and the CoM and treated it positive if the rotation is counter-clockwise with respect to axial direction of forward flight.

The flies were hit by the gust at mean axial location of X_0_ = 3.7 ± 0.38 D_0_ from the exit of the nozzle, and their lateral and vertical positions of CoM just before being hit by gust were at most 2L away from the centre of the gust (Figure 3A). If the flies were to the left side of the gust, they continued to fly in the same side even after being hit by the gust. Similarly, they flew downward after the encounter with gust, indicating a possible loss in lift forces. Further, the flies didn’t recover their initial vertical and lateral position after gust in any of the cases.

**Figure 3:**
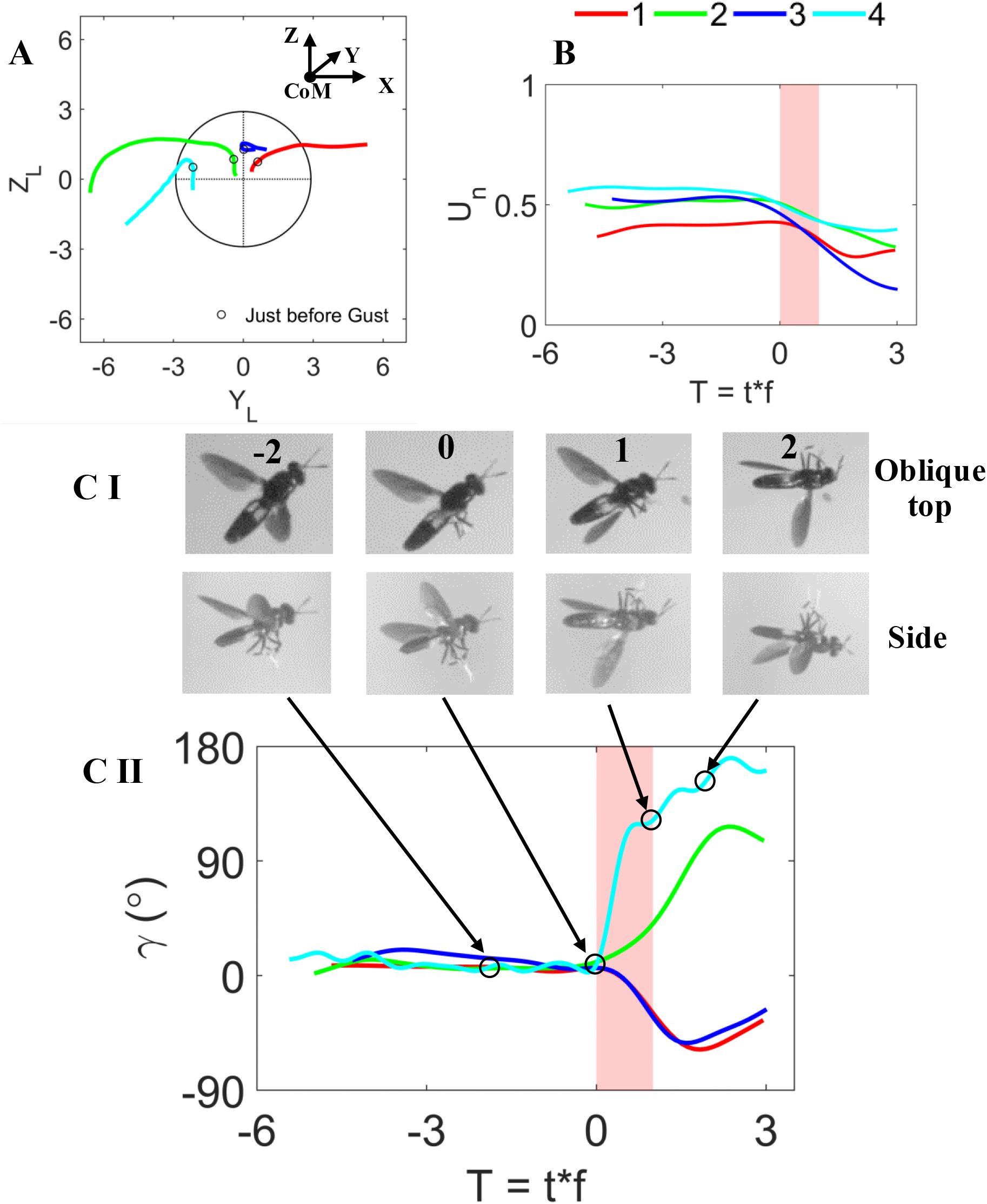
Changes in body kinematics of flies due to the ring in 4 different trials. (A) Trajectory in Y-Z plane normalized by the average body length (L_avg_) of flies. CoM is centre of mass. The mean axial distance where the gust hits the insects is X_0_ = 3.7 ± 0.38D_0_. Black circle indicates the front view of the vortex ring, and the intersection of vertical and horizontal dashed lines is the centre of the vortex ring. Coloured lines are the trajectories of flies for each trial represented by 1 −4, and open circles on each curve denote the position of the flies just before they were hit by the ring. (B) Normalized forward speed versus non-dimensional wing beat, the former normalized with the average speed of the flies before being hit by the ring and the latter is non-dimensionalized with the wing beat frequency. T=0 indicate the time instance just before flies were just hit by the ring. Time period of gust is indicated in vertical pink strip. (C-I) Oblique top and the corresponding side views of flight sequences for trial 4 at different time instances showing distinct change in body roll angle. Number on the top row indicates the time instance of fly with respect to gust in terms of wing beat. (C-II) Change in body roll angle plotted against the wingbeat. Open circles on blue curve (trial 4) denote the time instances of the fly shown in (C-I).

Before being hit by the ring, the forward speed of the flies was near-constant in each case. However, the gust resulted in a decrement in their forward speed by as much as ~70% maximum and ~30% on average (Figure 3B). Similarly, the flies had near-zero body roll angle (< 15°) before they come in contact with the ring, but it changed by as much as ~160° due to gust (Figure 3C). The flies also responded to the ring by varying other body angles and wing stroke angle as well, the detailed results of which will be discussed in the companion paper. It is, thus, evident from the observed changes in the flight trajectory, speed and body orientation, that the vortex ring can be used as a gust to perturb flies.

## CONCLUSION

We here presented a method of gust generation in form of a discrete vortex ring. The ring was generated by an impulsive motion of the diaphragm of an electronic speaker. In contrast to the conventional methods of gust generation – some of which are either not realistic or are difficult to characterize, the flow physics of this perturbation method is well understood, and the flow properties are highly controllable. Besides, high repeatability and reproducibility, low cost and simple mechanical design are the added advantages of this method. Furthermore, this method, in contrast to piston-cylinder arrangement of vortex ring generation, precludes the formation of any piston and secondary vortex. We have demonstrated its application for studying the impact of gust on insect flight. Further, the theoretical estimation, based on basic conservation equations, predicts the flow properties of the ring very well. The application of the vortex ring as a gust is not only limited to insects but could potentially be extended to study its impact on birds, bats, fishes and micro-aerial vehicles (MAVs) in air and on deep water sea plants and their growth, fish and underwater autonomous vehicles in water.

