## Supplemental files for "Design and development of a vortex ring generator to study the impact of the ring as a gust"

**Supplementary information**

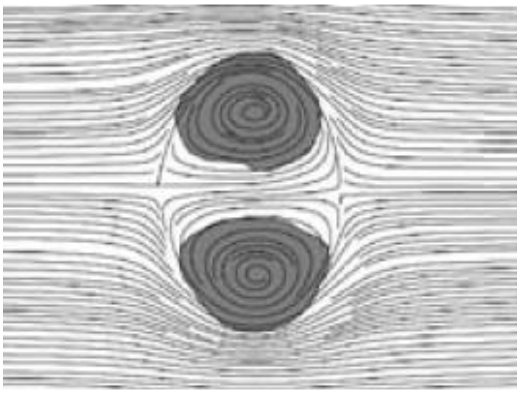

Figure S1: Instantaneous streamlines and vorticity patch at some time after the generation of vortex ring with respect to frame moving with it, taken from Dabiri and Gharib (2004).

*Table-S1: Physical measurement of five Styrofoam beads used for flow characterization*

| Diameter (m) x $10^{-3}$ | | Mass (kg) x $10^{-7}$ | Density ($Kg/m^3$) |
|---|---|---|---|
| | 6.527 | 8.7 | 5.98 |
| | 6.150 | 9.5 | 7.80 |
| | 6.653 | 6.6 | 4.28 |
| | 5.856 | 8.3 | 7.89 |
| | 6.091 | 7.9 | 6.67 |
| Mean | 6.26 | 8.2 | 6.52 |

### Velocity field of vortex ring

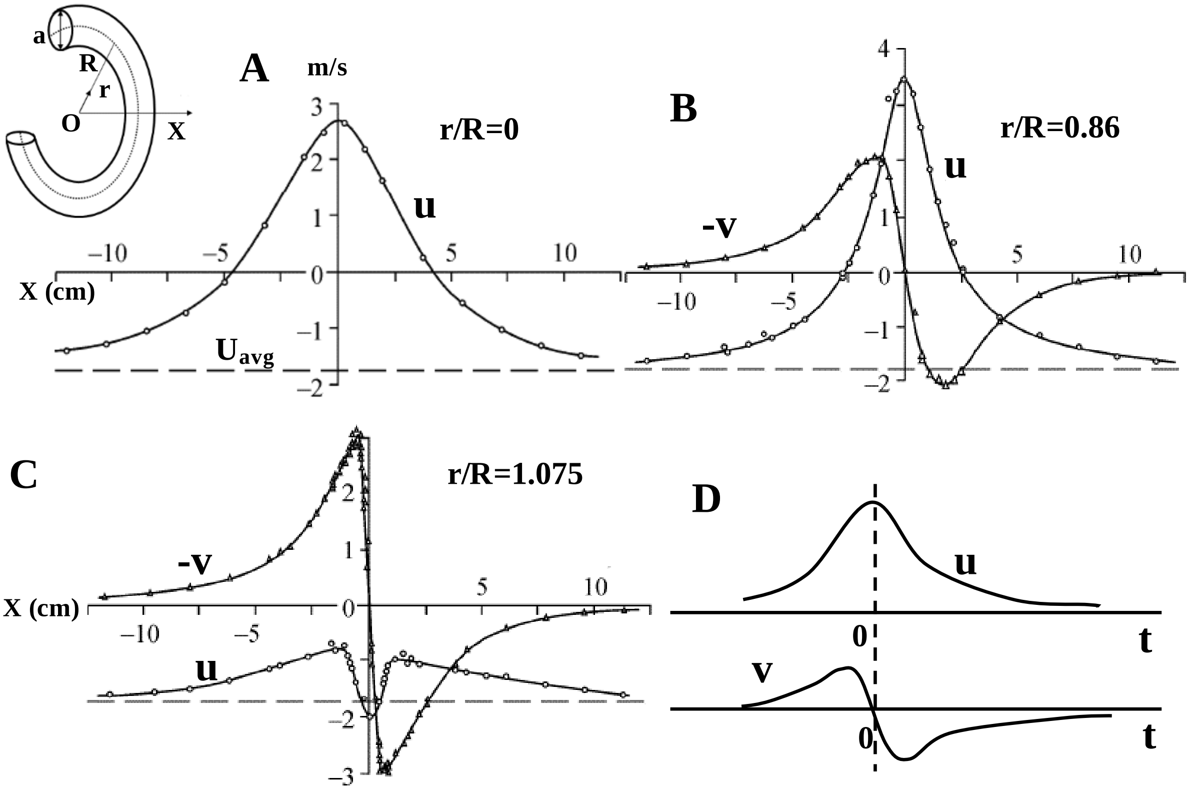


*Figure S2: Velocity field of a vortex ring in a frame moving with the ring. Vortex ring dimension with core is shown in the inset. X and r are axial and radial directions respectively and O is the centre of the ring in frontal plane and R and a are the radius of the ring and its core respectively. Axial (u) and radial (v) velocity distribution along X at the centre of the ring (r = 0) (A), at r = 0.86R (B) and at r = 1.075R (C) from the centre of the ring. Radial velocity is zero at the centre and it increases away from the centre. Axial velocity decreases greatly near the core (C). Note in (A, B and C), Re ($U_{avg}R/\upsilon$)=4.54 x $10^3$, solid lines are cubic splines fitted to the experimental data, and dashed lines denote the average translational velocity of the ring. Figures are adapted from Akhmetov (2009). Dashed line, if considered X-axis, would result in an axial velocity with respect to fixed laboratory frame, while radial velocity is invariant to such coordinate transformation. (D) Representative velocity distribution with time at any radial location, except at the centre where the radial velocity is zero.*

This section is intended to present some basic ideas about the velocity field of the vortex ring. While the vortex ring has an average translational velocity, the fluid associated with it has axial and radial velocity as well, and these velocities are different from their translational velocity (Figure S2 A-C). The ring has only the axial velocity at the centre of the ring in the frontal plane while the radial velocity is zero at that point (Figure S2 A). The magnitude of both velocity components increases away from the centre of the ring (Figure S2 B,C), reaching maximum value just before the core region and decreasing to a minimum beyond that. Indeed, the distance between the maximum and the minimum value of axial (or radial) velocity component gives the measure of the diameter of the core (Das et al. 2017). Velocity field of these components vary in similar fashion with time (Figure S2 D).
